# A new nanoDSF approach to anti-tubulin compounds screening revealed novel MTAs among approved drugs

**DOI:** 10.1101/2024.12.26.630395

**Authors:** Viktoriia E Baksheeva, Romain La Rocca, Ludovic Leloup, Aurelie Souberan, Raphael Bergès, Diane Allegro, Carine Derviaux, Eddy Pasquier, Philippe Roche, Xavier Morelli, François Devred, Aurélie Tchoghandjian, Emeline Tabouret, Andrey V Golovin, Philipp O Tsvetkov

## Abstract

Microtubule Targeting Agents (MTAs) constitute a vital category of tubulin-binding compounds, deployed across anticancer therapies. Despite the array of MTA drugs developed by pharmaceutical entities, the quest for novel efficacious molecules continues unabated. We unveil an innovative *in vitro* MTA screening methodology employing nano differential scanning fluorimetry (nanoDSF), presenting distinct advantages over known assays. This novel approach not only assesses compound-tubulin binding but also quantitatively analyzes their impact on tubulin polymerization. Proposed nanoDSF assay was rigorously validated using the Prestwick Chemical Library, which encompasses 1,520 approved compounds, successfully identifying all previously known MTAs. Furthermore, this screening has unearthed potential anti-tubulin agents among drugs currently utilized for non-related medical conditions, offering insights into their mechanisms of action in inhibiting cancer cell proliferation and/or inducing cytotoxicity. These discoveries herald new opportunities for drug repositioning involving the newly identified MTAs and substantially streamline the process of screening extensive chemical libraries for MTAs featuring novel chemical structures.

## Introduction

Microtubules (MT), polymeric structures composed of tubulin heterodimers, possess the dynamic ability to elongate or shorten (Fig.1A) in response to environmental shifts. The dynamic behavior of MTs is essential for cell motility and division, playing key roles in cytoskeletal rearrangement during cell migration and the accurate positioning of chromosomes during mitosis. This necessitates a sophisticated level of control over MT dynamics. To achieve such precise regulation, cells have developed a complex network of microtubule-associated proteins (MAPs) that meticulously orchestrate the dynamics of MTs^1^. Any disruption in this fine-tuned process can adversely affect cell migration, block mitosis, and thus potentially be fatal for the cell. This explains why MT dynamics have become the focus of a wide range of pharmacological agents.

**Figure 1.**
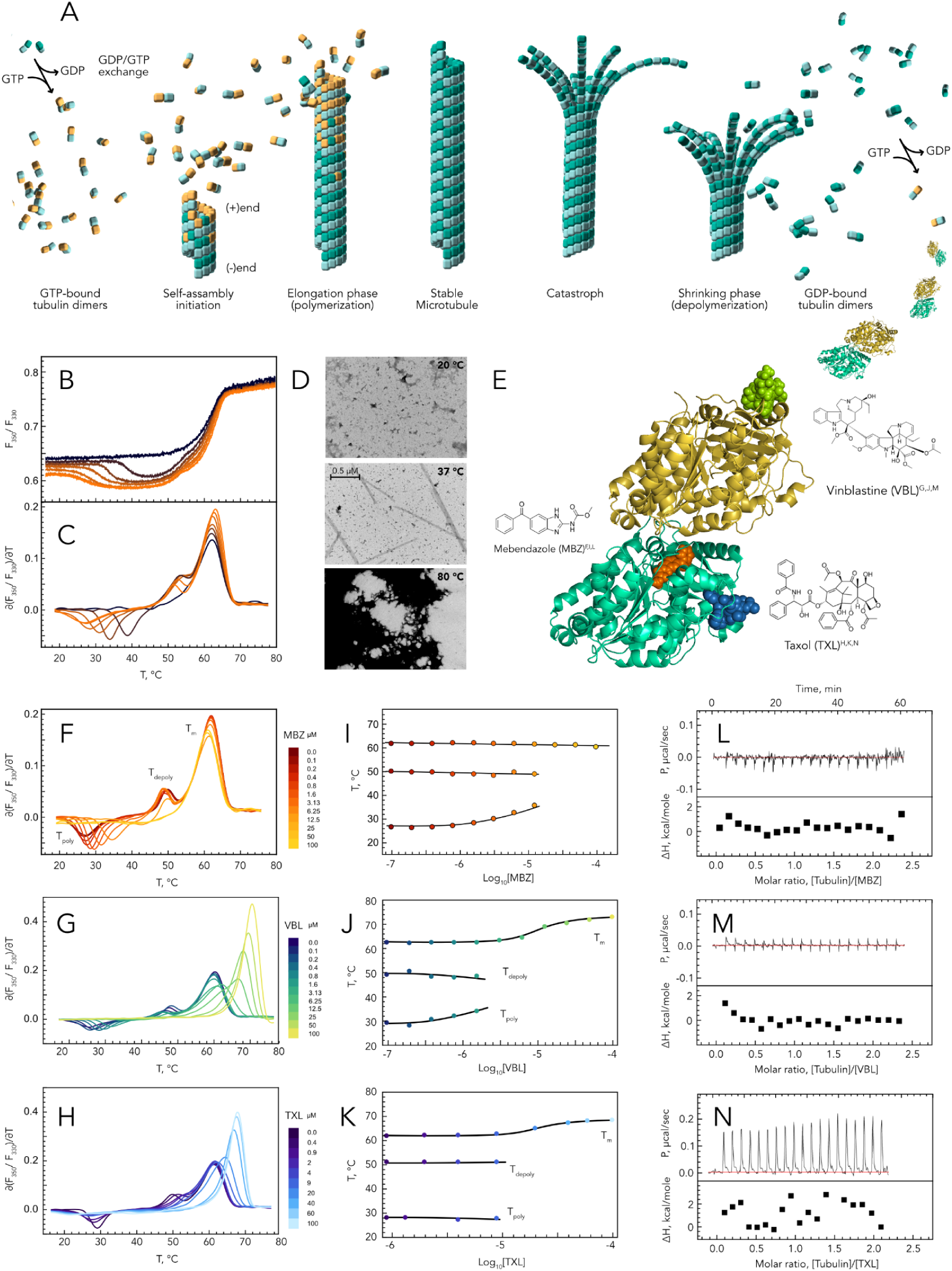
Monitoring tubulin polymerization with nanoDSF. (A) Different stages of tubulin polymerization in microtubules. (B,C) Polymerization and denaturation of tubulin at different concentrations followed by nanoDSF from 20 to 75°C. (D) TEM of tubulin at different temperatures. (E) Structures of mebendazole (MBZ), vinblastine (VBL), and taxol (TXL) molecules and its complex with tubulin heterodimer. (F,G,H) Polymerization and denaturation of tubulin in the presence of different concentrations of MBZ, VBL, and TXL respectively followed by nanoDSF. (I,J,K) Dependance of the temperatures of polymerization (T_poly_), depolymerization (T_depoly_) and denaturation (T_m_) of tubulin from concentration of MBZ, VBL, and TXL respectively. (L,M,N) Isothermal titration calorimetry curves of tubulin titration by MBZ, VBL, and TXL respectively in polymerization buffer.

Microtubule targeting agents (MTAs) is a class of tubulin-binding compounds that is used not only for cancer treatment but also as anthelmintic, antibacterial and antifungal drugs. Moreover, recently MTA administration was proposed as a new strategy for the therapy of neurodegenerative diseases ^2,3^. MTA interaction with tubulin significantly impacts MT dynamic, thus perturbing vital cellular processes. MTAs differ by their binding sites on tubulin ^4,5^ (Fig. 1E) and are grouped in two classes by their ability to induce or inhibit MT formation. The first MTAs with anti-cancer activity were originally obtained from plants ^6^ and were subsequently subjected to chemical modifications to yield new compounds with the superior anti-cancer properties. While pharmaceutical companies have developed various MTA anticancer drugs, the demand for new molecules remains high, driving researchers to explore additional natural sources of MTAs^7^. These new compounds should exhibit increased specificity and broader efficacy across different tumor types and should be capable of overcoming drug resistance observed in certain tumors, which may arise from MTA interplay with MAPs^8,9^. Unfortunately, the strategy of chemical modifications of existing drugs does not allow to create MTAs with significantly different characteristics since they all possess similar scaffolds. Thus, to identify MTAs with potentially superior anticancer potential, it is necessary to screen large and diverse libraries of chemical compounds.

There are several molecular screening assays based on different biophysical methods that allow to identify interaction between a target protein and potential binders. Among them thermal shift assay (TSA) became widely used for early-stage drug discovery in the last years because it is accessible (RT-PCR equipment is sufficient), high throughput (adapted for 96, 384 and 1,536 plaques) with a low material consumption^10^. This technique facilitates the assessment of protein thermostability (melting or denaturation temperature, T_m_) in the presence of screening compounds, thereby providing evidence of their interaction. However, like many molecular assays, it generates a certain number of false positive and false negative results. This phenomenon arises because, first, not all interactions between the protein and compounds lead to a significant change in protein thermostability, as detected by TSA, and second, not every interaction impacts the biological process regulated by the protein, which is the intended target of new drug therapies. This is especially relevant for tubulin, which possesses multiple ligand-binding sites. Consequently, there is no direct correlation between the extent of tubulin stabilization by a compound and its efficacy as a polymerization inhibitor or promoter. Thus, molecular screening assays that not only demonstrate tubulin-compound interactions but also elucidate the effect of the compound on tubulin polymerization (the specific process targeted) are essential for the efficient discovery of MTAs.

To address these challenges, we introduce an *in vitro* functional MTA screening methodology utilizing nanoDSF. Unlike RT-PCR based assays, nanoDSF does not require fluorescent dyes, as it monitors the intrinsic fluorescence of proteins. Remarkably, this approach enables a dual readout, allowing for simultaneous detection of both tubulin stabilization by the compound and its effect on tubulin polymerization. We applied the nanoDSF screening assay to the Prestwick Chemical Library (PCL), which comprises 1,520 approved compounds. This application not only validated the assay with known MTAs contained within the library but also led to the identification of approximately 100 molecules demonstrating MTA activity. Finally, we investigated the anti-cancer activity of the identified hits on glioblastoma cell lines (GBM) and tested some of them on patient-derived tumoroids, demonstrating the integration of our screening approach across the entire drug development cycle, from initial screening to patient application.

## Results

### Monitoring tubulin polymerization using nanoDSF

To monitor tubulin polymerization, we used a nanoDSF instrument from NanoTemper Technologies, which enables tracking of the protein’s intrinsic fluorescence at 330 nm and 350 nm across a temperature range of 15°C to 95°C. The ratio of these two signals (F_330_/F_350_) reflects the exposure of the protein’s aromatic amino acids to the solvent, allowing for the observation of protein denaturation upon heating the sample. We have noticed that the dimerization interface of tubulin contains several aromatic amino acids, potentially enabling the use of nanoDSF not only to monitor tubulin denaturation but also to observe the formation of MTs. To test this hypothesis, we subjected tubulin samples at varying concentrations to heating using nanoDSF in a polymerization buffer, wherein tubulin has the capability to polymerize upon reaching physiological temperatures (Fig. 1B,C). At a low subcritical concentration of tubulin, we detected only a single transition around 62°C, indicative of tubulin denaturation denoted further as T_m_. Consistently, at higher concentrations of tubulin, two additional transitions emerged on the thermogram. The first transition, occurring at approximately 28°C, displayed a signal opposite to that of tubulin denaturation, suggesting that aromatic amino acids were concealed from the solvent rather than exposed. This event most probably corresponds to tubulin polymerization. The subsequent transition, occurring near 55°C, matched the magnitude of signal change of the 28°C transition but was opposite in direction, strongly suggesting it correlates with the depolymerization of MTs. The polymer state of the tubulin under nanoDSF experimental conditions was confirmed using TEM microscopy at different temperatures (Fig. 1D), thus validating nanoDSF as a potential tool to study the impact of compounds on tubulin polymerization. Henceforth, we will denote the temperatures corresponding to the minima and maxima of the first derivative of the (F_330_/F_350_) ratio as the apparent temperatures of tubulin polymerization (T_poly_) and depolymerization (T_depoly_), respectively.

Given that tubulin has multiple binding sites influencing its polymerization, we evaluated the interaction of tubulin with three MTAs - Vinblastine (VBL), Taxol (TXL), and Mebendazole (MBZ) - each binding to distinct sites (Fig. 1E). This was done to assess the capability of the nanoDSF assay to detect interactions between tubulin and MTAs. Therefore, tubulin samples, in the presence of increasing concentrations of MBZ, VBL, and TXL, were subjected to heating from 15°C to 75°C using the nanoDSF instrument. This induced markedly distinct alterations in tubulin polymerization and denaturation profiles (Figs. F, G, H). A gradual increase in MBZ concentration resulted in a notable shift of T_poly_ to higher temperatures without affecting T_depoly_ and T_m_ (Fig. 1F, I), until reaching a concentration of 100µM. At this concentration, MBZ entirely inhibited MT formation, resulting in the elimination of the first two transitions (Fig. 1F, yellow curve). Contrary to MBZ, VBL is able not only to inhibit tubulin polymerization in sub stoichiometric amounts, but also to decrease the temperature of depolymerization (Fig. 1 G,J). Moreover, saturation of tubulin with VBL leads to significant stabilization of tubulin structure. TXL, known to promote tubulin polymerization in contrast to MBZ and VBL, causes T_poly_ and T_depoly_ to shift in opposite directions (refer to Fig. 1H, K). Like VBL, TXL stabilizes the tubulin structure; however, distinctions between TXL and VBL’s interactions with tubulin are observable. Specifically, a detailed examination of tubulin denaturation peaks with increasing concentrations of VBL reveals a sequence where initially, the peak’s magnitude decreases, followed by the peak becoming asymmetric, and ultimately, it increases in amplitude and shifts to higher temperatures. In contrast, TXL leads to a gradual increase in both the amplitude and T_m_ of tubulin’s symmetric denaturation peak. These findings align well with the known locations of TXL and VBL binding sites on tubulin, which are responsible for the distinct dynamics of compound exchange between the free and bound states, thereby differentially affecting tubulin denaturation. Ultimately, the apparent affinity constants of compounds could be independently estimated by analyzing both the T_m_ shift and the alterations in fluorescence signal at 15°C, which is particularly advantageous in cases where conventional reference methods, like Isothermal Titration Calorimetry (ITC), are ineffective under polymerization conditions (Fig. 1L,M,N).

### Application of nanoDSF for MTA screening

Thus, tracking temperature-induced tubulin polymerization with nanoDSF enables the detection of shifts in the polymerization temperature (ΔT_poly_) across a broad concentration range with high sensitivity. This approach facilitates the qualitative assessment of MTAs’ effects on tubulin polymerization, allowing for their comparative evaluation based on this criterion. Additionally, further heating reveals the impact of MTAs on tubulin’s thermostability (ΔT_m_). By measuring these two distinct parameters - the first directly related to the tubulin function targeted by MTAs and the second reflecting the structural influence of MTAs on tubulin - this approach emerges as highly promising for MTA screening. Advanced nanoDSF instruments, such as the automated Prometheus NT.Plex, are equipped to conduct high-throughput screening of chemical libraries for compounds targeting tubulin.

To evaluate the efficacy of our novel MTA screening methodology, we applied it, as a proof of concept, to the PCL, which comprises 1,520 approved drugs (Fig. 2A-C). We observed that in the presence of 20 compounds (1.3% of PCL), tubulin exhibited no polymerization, suggesting either a complete inhibition of MT formation or the initiation of MT formation at temperatures below 15°C. These compounds are henceforth categorized as strong hits. Among these, nine are already known as MTAs (Table 1), three are suspected of having MTA activity (Auranofin, Ebselen, and Riboflavin), and eight (Aprepitant, Benzarone, Benzbromarone, Benziodarone, Bithionol, Hexachlorophene, Nifedipine, and Nisoldipine) are newly identified as exhibiting MTA activity, previously unreported. For the remaining 1,500 drugs, tubulin polymerization occurred, enabling their classification based on two metrics: ΔT_poly_ and ΔT_m_. The initial findings, along with 1D and 2D distributions of these metrics, are depicted in Figures 2B,C. Through statistical analysis of the results (Fig. 2D), compounds causing a ΔT_poly_ shift greater than 2°C were designated as true MTAs. Those with a ΔT_poly_ shift as low as 1°C are also considered potential MTAs, termed weak hits, meriting further investigation.

**Figure 2.**
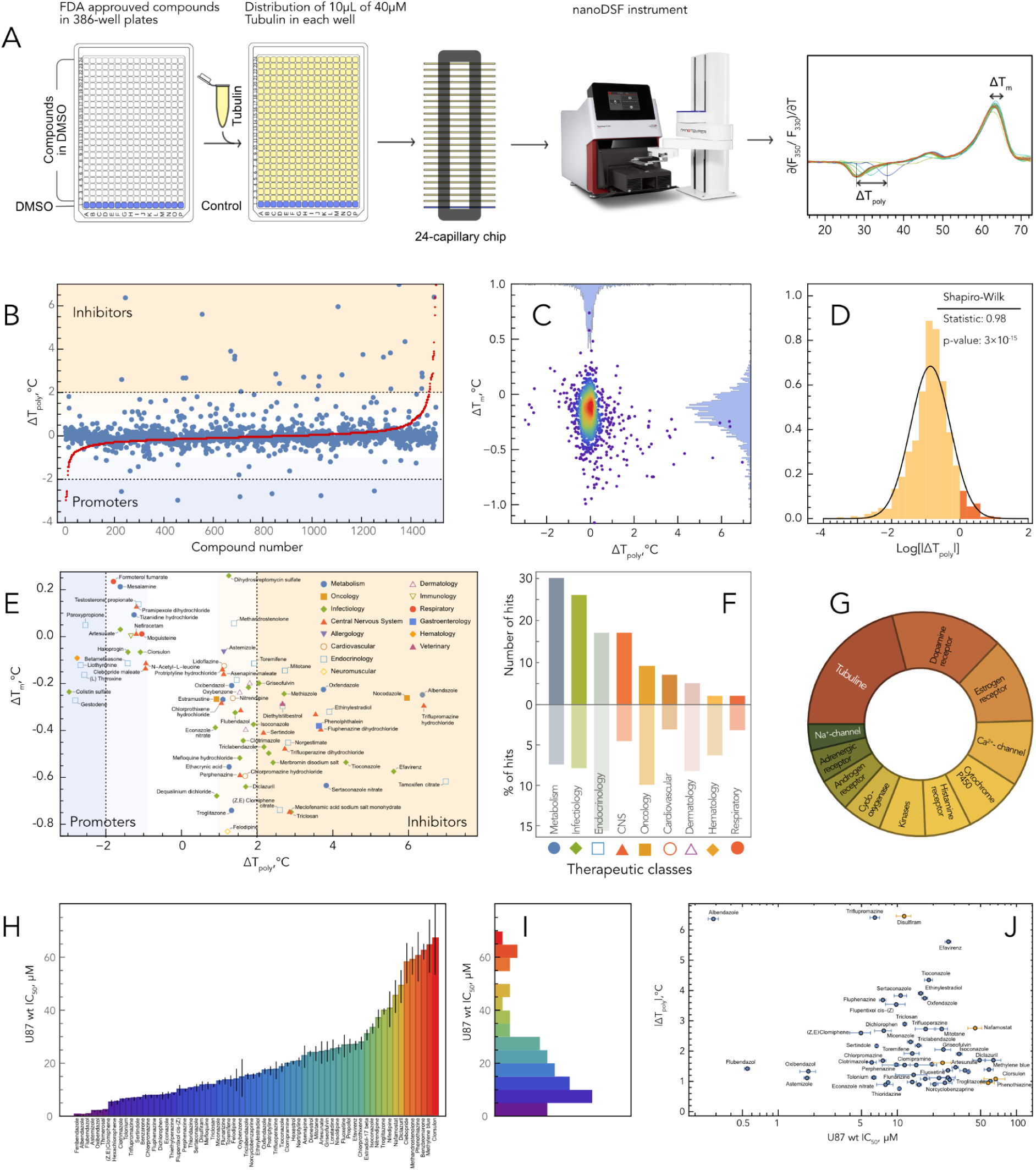
nanoDSF screening of PCL of 1,520 approved compounds. (A) nanoDSF screening workflow. (B) Results of PCL screening. (C) Distributions of ΔT_poly_ and ΔT_m_ values represented as 1D histograms and 2D density plot. (D) Distributions of Log_10_|ΔT_poly_| and its fitting. (E) 2D distributions of ΔT_poly_ and ΔT_m_ values of hits with their therapeutic classes. (F) Distribution of hits in therapeutic classes. (G) Known primary targets of hits. (H, II) IC_50_ of hits for U87MG cancer cell line and its distribution. (J) 2D distributions of absolute value of |ΔT_poly_| and IC_50_ of hits (orange points have negative ΔT_poly_).

**Table 1.** The list of drugs from PCL with the most important effect on tubulin polymerization.

| Name | Therapeutic class | Targets | Impact on microtubules | Cancer treatment |
| --- | --- | --- | --- | --- |
| **Known microtubule targeting agents** |  |  |  |  |
| **Colchicine** | Metabolism | **Tubulin** | Prevents microtubule assembly and thereby disrupts inflammasome activation.<sup>28</sup> | Derivatives regarded as potential chemotherapy drugs.<sup>76</sup> |
| **Docetaxel** | Oncology | **Tubulin** | Disrupts normal microtubule dynamics and thereby stops cell division.<sup>77</sup> | Chemotherapy agent utilized in various cancers.<sup>78</sup> |
| **Fenbendazole** | Infectiology<br>Metabolism | **Tubulin** | Moderate affinity to mammalian tubulin.<sup>40</sup> | Moderate antineoplastic activity.<sup>40</sup> |
| **Mebendazole** | Infectiology<br>Metabolism | **Tubulin** | Selectively inhibits tubulin polymerization via interaction with colchicine-binding site of β-tubulin.<sup>79</sup> | Repositioned as prospective anti-cancer agent.<sup>39</sup> |
| **Paclitaxel** | Oncology | **Tubulin** | Stabilizes the microtubule polymer and protects it from disassembly.<sup>80</sup> | Anti-cancer drug.<sup>81</sup> |
| **Podophyllotoxin** | Metabolism | **Tubulin** | Prevents polymerization of tubulin by binding to colchicine site.<sup>82</sup> | Anti-tumor, derivatives applied in chemotherapy.<sup>83</sup> |
| **Dienestrol** | Endocrinology | ERα, ERβ | Inhibits microtubule assembly *in vitro* by binding to site analogous to colchicine site.<sup>17</sup> | Carcinogenic.<sup>84</sup> |
| **Hexestrol** | Endocrinology<br>Oncology | ERα, ERβ | Inhibits microtubule assembly *in vitro* by binding to site analogous to colchicine site.<sup>17</sup> | Carcinogenic.<sup>85</sup> |
| **Thiomersal** | Infectiology | InsP<sub>3</sub>R | Inhibits tubulin polymerization *in vitro*.<sup>18</sup> | Induces apoptosis in some cancer cell lines.<sup>86</sup> |
| **Suspected for MTA activity** |  |  |  |  |
| **Auranofin** | Metabolism | NF-κB kinase $\beta$; PRDX5 | Inhibits phagocytosis which could be linked to microtubule modulation.<sup>20</sup> | Cytotoxic to mutant p53 cancer cells.<sup>37</sup> |
| **Ebselen** | Metabolism<br>CNS | AChE, SEH | In high doses disrupts microtubules in cells.<sup>23</sup> | Suppresses cancer cell growth.<sup>24</sup> |
| **Riboflavin** | Metabolism<br>Ophthalmology | FMN | Rescues cytoskeletal alterations in patients with RTD.<sup>25</sup> | Potential adjuvant in chemoradiotherapy.<sup>26</sup> |
| **Newly detected MTAs** |  |  |  |  |
| **Aprepitant** | Metabolism | NK 1 receptor | Unknown | Potential anti-tumor agent.<sup>31</sup> |
| **Benzarone** | Rheumatology | Nonpurine XO; SLC22A12; EYA3 | Unknown | Inhibits tumor growth in animal model.<sup>27</sup> |
| **Benzbromarone** | Cardiovascular | Uric acid uptake | Unknown | Predicted as therapeutic drug for LUAD<sup>35</sup> |
| **Benziodarone** | Cardiovascular | Uric acid uptake | Unknown | Not used |
| **Bithionol** | Dermatology | ADCY1 | Unknown | Synergistic with paclitaxel in ovarian cancer.<sup>33</sup> |
| **Hexachlorophene** | Infectiology | G6PDH, SHP2 | Unknown | Suppresses proliferation in NSCLC model.<sup>33</sup> |
| **Nifedipine** | Cardiovascular | Cytochrome P450 3A4 | Unknown | Reverses drug resistance of cancer cells.<sup>34</sup> |
| **Nisoldipine** | Cardiovascular | DHP channel | Unknown | Not used |

Among both strong and weak hits, we discovered that 30 compounds are classified within the therapeutic category targeting metabolism, 26 are utilized in infectious diseases, 17 in endocrinology, and 17 in the treatment of central nervous system disorders (see Figs. 2E,F). When examining the therapeutic effects of the drugs that influence tubulin polymerization, we identified 27 compounds with antifungal properties, 19 with antineoplastic effects, 16 with antibacterial activity, and 9 each with anti-inflammatory, antipsychotic, and anthelmintic effects. As expected, the main known target of those hits was tubulin, still for more than 75% of hits has other “main” target protein (Fig. 2G). Therefore, tubulin should also be considered a significant target for these compounds, which could explain the molecular mechanism of action of some compounds, or the side effects associated with the clinical use of these molecules.

To assess whether compounds with MTA activity also demonstrate cytotoxic effects, cell survival assays were conducted on the human U87MG glioblastoma cell lines at varying concentrations of the identified compounds. Our analysis revealed that for approximately 50% of these compounds (54 molecules), the IC_50_ value was less than 40µM, while for about 20% (19 compounds), it was under 10µM (Fig. 2H-J). The IC_50_ values showed a modest negative correlation with the change in tubulin polymerization temperature (ΔT_poly_) (Fig. 2K).

### Structure-activity relationship of some newly identified MTAs

Furthermore, we analyzed the structural similarity among all hit compounds. To achieve this, we initially computed a structure similarity matrix detailing the pairwise distances between compounds, utilizing ‘Morgan Connectivity’ fingerprints. Subsequently, we derived a cluster hierarchy based on this matrix (Fig. 3A) and reorganized the structure similarity matrix in accordance with the identified clustering, excluding compounds with minimal structural resemblance (Fig. 3B, hits with small structural similarity are listed in Table S8). This process enabled us to identify and better visualize several clusters of molecules with analogous structures within the hits (Fig. 3A-B). Thus, alongside various small clusters that include both previously identified and novel MTAs (Fig.3A,B, Tables S3-S7), two prominent clusters are highlighted, composed of established microtubule inhibitors: carbendazim (Fig. 3D, Table S2) and phenothiazine derivatives (Fig. 3E, Table S1). In the last cluster we identified two pairs of molecules, perphenazine (PPZ) and fluphenazine (FPh), as well as chlorpromazine (CPZ) and triflupromazine (TFZ), wherein the substitution of a chlorine atom (-Cl) at the 2^nd^ position of the phenothiazine scaffold with a trifluoromethyl group (-CF_3_) (see Fig. 3D) lead to significant increase in ΔT_poly_. To gain deeper understanding of this phenomenon, we employed funnel metadynamics to simulate the docking of these four molecules, along with colchicine as a control, into the colchicine binding site of α-tubulin — also recognized as the binding site for phenothiazine derivatives ^11^. The docking of colchicine to α-tubulin resulted in a center-of-mass position that was consistent with X-ray crystallographic data (data not shown). Next, a comparative analysis of the two pairs of compounds showed that the introduction of trifluoromethyl groups generally altered both the position and affinity of the molecules (Fig. 3F,G). In the CPZ-TFZ pair, the difference was most pronounced, with the trifluoromethyl group penetrating deep into the protein cavity and "dragging" the entire molecule with it. In the PPZ-FPh pair, the trifluoromethyl group also played a key role in forming effective contacts within the hydrophobic region of the β-sheet near the colchicine binding site. According to our calculations, the affinity of FPh was significantly higher than that of PPZ. Comparison of TFZ and FPh suggests that their inhibition efficiencies arise from different mechanisms: while FPh exhibits high affinity, TFZ binding leads to substantial rearrangements in the interfacial interactions between α- and β-tubulin subunits.

**Figure 3.**
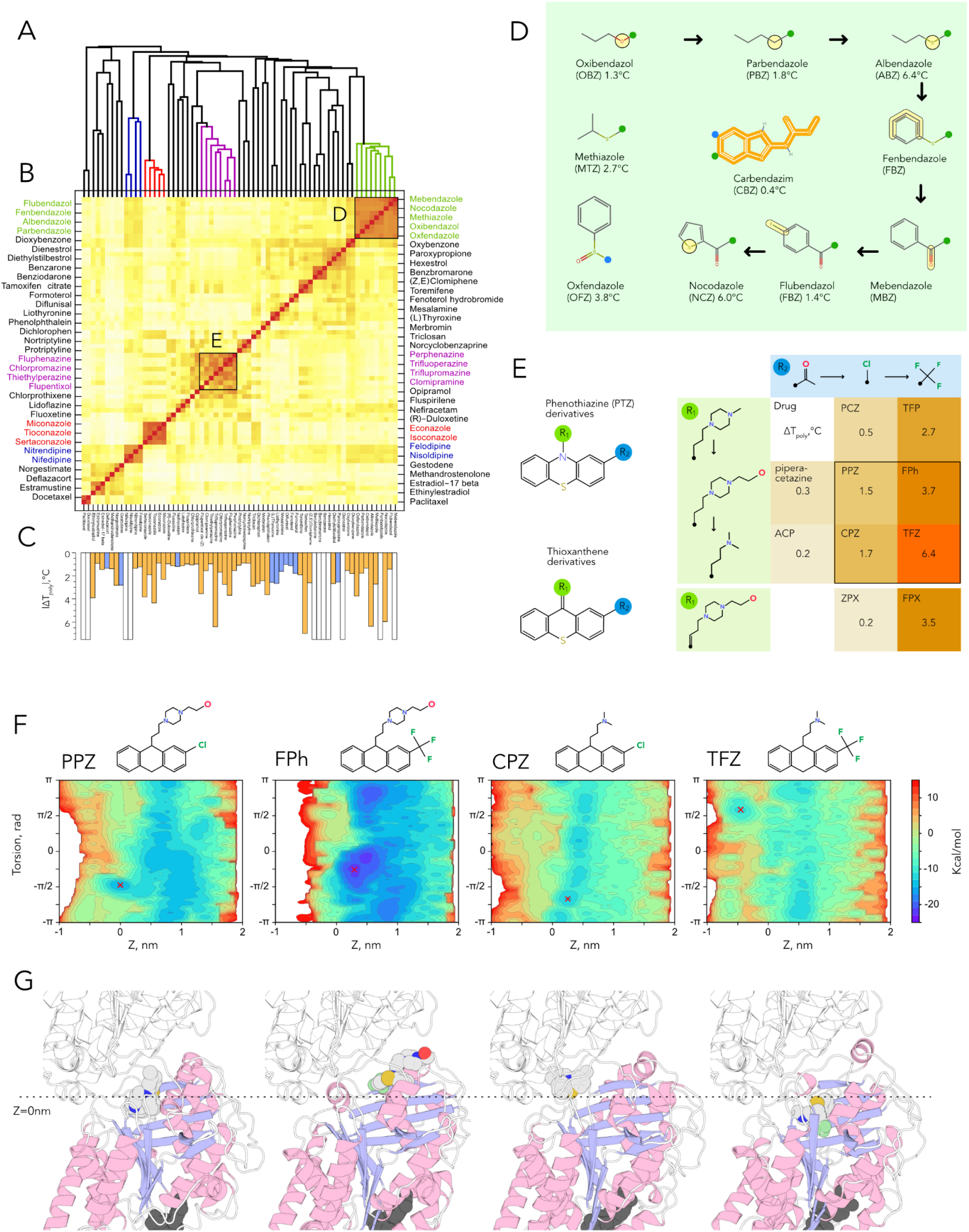
Structure-activity relationship of hits and funnel metadynamics simulation results. (A) Hierarchical dendrogram representing chemical clusters of hits. (B) Sorted chemical similarity matrix for hit compounds that have at least one similar compound among the hits. (C) Absolute value of ΔT_poly_ of hits: polymerisation inhibitors are shown in orange and promoters in blue bars. Strong hits that completely inhibit tubulin polymerization are shown in white bars. (D) Structures and ΔT_poly_ of CBZ derivatives. Each next modification is highlighted in light yellow. (E) Structures and ΔT_poly_ of PTZ derivatives. (F) Funnel metadynamics simulation of the interactions between compounds PPZ, FPh, CPZ, and TFZ upon binding with the α-tubulin subunit. The 2D plot represents the free energy profile of compound binding, with coordinates as follows: the x-axis indicates the distance from the center of mass of the CLH (as determined by X-ray data, zero value) to the center of mass of the compound, while the y-axis shows the torsion angle representing the arbitrary rotation of the compound molecule relative to tubulin within the interaction plane. Stable binding modes are marked with a red cross. (G) Visualization of the binding modes of compounds PPZ, FPh, CPZ, and TFZ with corresponding α-tubulin conformations denoted by red cross minima. The protein molecules are shown in cartoon mode, colored in blue and light pink, while the compound molecules are displayed as spheres with carbon atoms in gray, chlorine atoms in yellow, and fluorine atoms in light green. The dashed line represents the position of the CLH center of mass.

### Toward potential repurposing of perphenazine derivatives for GBM therapy

Since phenothiazine derivatives are known to cross the blood-brain barrier, they may hold therapeutic potential for brain tumors with typically poor prognosis. Specifically, we focused on two pairs of phenothiazine derivatives that demonstrated the highest ΔT_poly_ and lowest IC_50_ values in GBM U87MG cell lines. Although inhibiting microtubule assembly is essential for the anticancer activity of MTAs, it alone is not sufficient for effective treatment. While we confirmed the activity of these compounds in U87MG glioma 2D cell lines, the 3D structure of tumors can hinder drug penetration, leading to variations in drug efficacy between 2D and 3D models. To address this, we extended our study to assess the effects of these four molecules on U87MG-derived spheroids, which better mimic the environment in tumors. Notably, unlike in 2D models, CPZ and TFZ exhibited higher IC_50_ values in spheroids (27.6±1.3 µM and 24.6±2.4 µM, respectively) compared to PPZ and FPh (17.9±1.7 µM and 17.0±0.8 µM) (Fig. 4A,B). This suggests that PPZ and FPh may have enhanced efficacy in a more complex cellular environment. This phenomenon may be attributed to the enhanced penetration capabilities into 3D cellular architecture of molecules that feature a 2-(4-Propyl-1-piperazinyl)ethanol group at the 10^th^ position on the phenothiazine scaffold (Fig. 3D). These results suggest that PPZ and FPh are the most promising candidates for repurposing as glioblastoma treatments.

**Figure 4.**
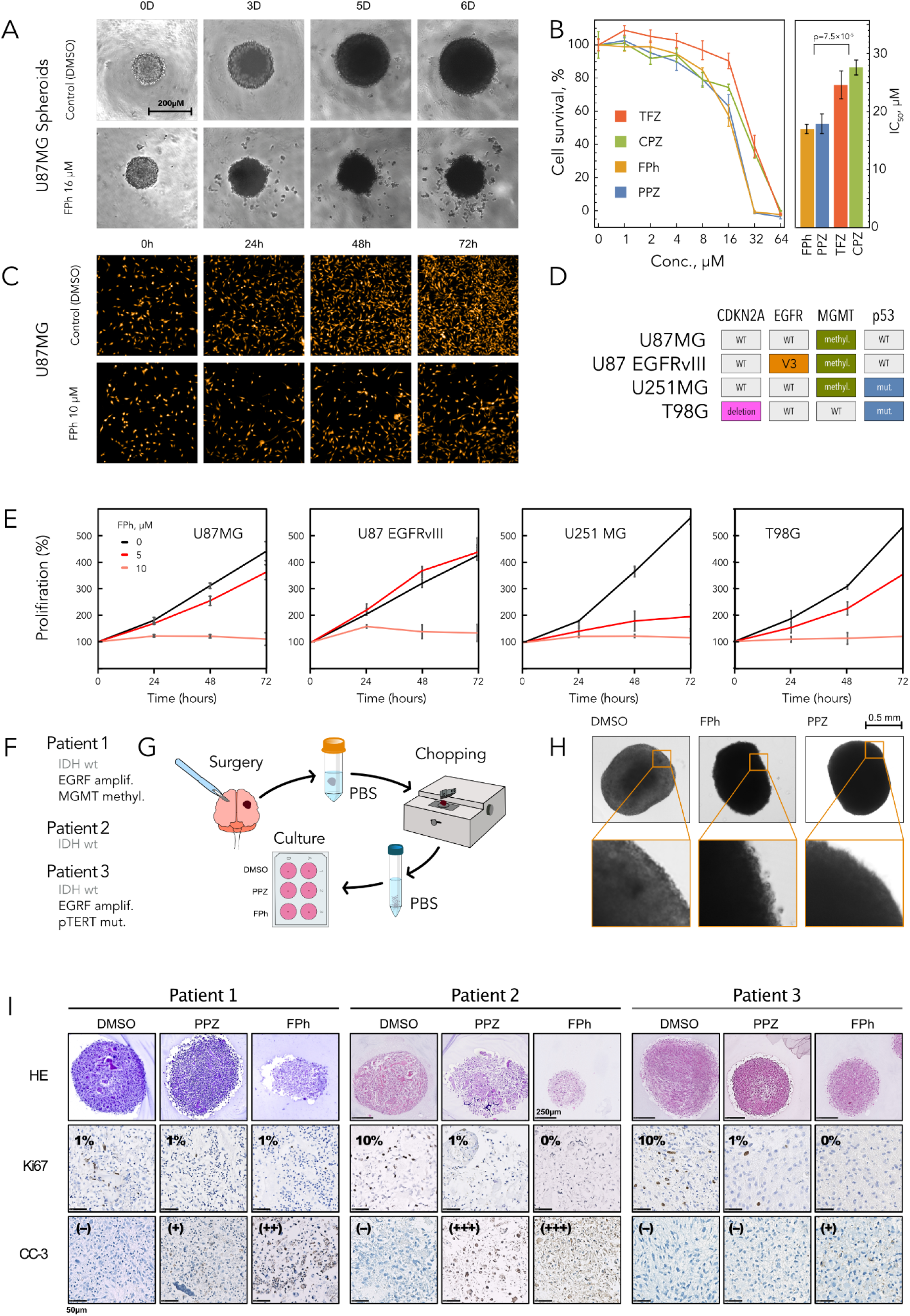
Cytotoxicity and antiproliferative effects of phenothiazine derivatives. (A) Representative images of U87MG spheroids treated with 16 μM FPh over a 6-day period, highlighting the impact on spheroid morphology. (B) Effects of increasing concentrations of TFZ, CPZ, PPZ, and FPh on U87MG spheroid cell viability. (C) Images from the Operetta High Content Imaging system illustrating the antiproliferative effects of FPh on U87MG cells in artificial-coloring. (D) Molecular alterations present in glioblastoma cell lines, reflecting the genetic landscape commonly associated with GBM. (E) Effects of escalating concentrations of FPh on the proliferation of different glioblastoma cell lines. (F) Molecular profiling of patient-derived tumors, showing key genetic alterations. (G) Summary of the workflow for processing patient samples, leading to tumoroid generation. Ten (n=10) tumoroids were treated per condition. (H) Tumoroids treated for 72 hours with vehicle (DMSO), 17 μM FPh, or PPZ, showing distinct changes in tumoroid structure. (I) Immunostaining of tumoroids treated with PPZ or FPh, showing reduced proliferation (Ki67 staining) and increased apoptosis (cleaved-caspase-3 staining).

We further evaluated the antiproliferative effects of the selected compounds in corresponding sublethal consentrations on four glioblastoma cell lines (U87MG, U87 EGFRvIII, U251MG, and T98G), which exhibit molecular alterations commonly found in GBM (Fig. 4D). Cell proliferation was monitored using the Perkin Elmer Operetta High Content Imaging System, with images captured every 24 hours (Fig. 4C). Our results showed that PPZ stimulated the proliferation of U87MG and U87 EGFRvIII cells while modestly reducing proliferation in U251MG and T98G cells by 30.5±3.4% and 15.0±3.0%, respectively (Fig. S1). In contrast, both CPZ and TFZ reduced proliferation in U87MG, U251MG, and T98G cells (Fig. S1), but neither was effective in inhibiting U87 EGFRvIII growth. Interestingly, CPZ even promoted proliferation in U87 EGFRvIII cells by 28.8±3.0% (Fig. S1). FPh, however, stood out as the only compound with significant antiproliferative effects across all four cell lines, reducing cell growth by at least 89% (Fig. 4E). These findings suggest that the EGFRvIII mutation substantially limits the efficacy of several treatments but highlights the therapeutic potential of FPh, which robustly inhibits proliferation regardless of molecular alterations.

Given these promising results, we chose to further investigate FPh therapeutic potential. PPZ was used as a control, while CPZ and TFZ were excluded due to their ineffectiveness against U87 EGFRvIII cells and higher IC_50_ values in spheroids. To achieve this, we used IDH-wildtype glioblastoma patient-derived tumoroids (Fig. 4F,G). These tumoroids are "tumors-in-a-dish" that preserve the histological and molecular characteristics of the original tumors, as well as their cellular heterogeneity^12^. FPh effects were tested on tumoroids derived from three patients and compared with the effects of PPZ and the vehicle (DMSO). After 72 hours of treatment, PPZ-treated tumoroids retained a rounded shape, similar to vehicle-treated controls. In contrast, FPh-treated tumoroids displayed irregular edges, indicating morphological changes (Fig. 4H). To quantify these changes and assess whether FPh impaired proliferation and induced apoptosis in the tumoroids, we performed Ki67 and cleaved-caspase-3 immunostainings. FPh induced apoptosis in all three patient-derived tumoroids and completely halted cell proliferation in two out of three cases (Fig. 4I). In comparison, PPZ induced apoptosis in two out of three tumoroids and decreased proliferation in two out of three, although to a lesser extent than FPh (Fig. 4I).

## Discussion

### New approach for MTA screening

Microtubule targeting agents (MTAs) are an important class of compounds widely used in the treatment of various diseases, including antifungal, antibacterial, antihelminthic, and antineoplastic therapies. Moreover, there is a growing body of evidence that MT stabilizing MTAs could be used for treatment of brain disorders ^2,13^. Despite MTAs being considered a relatively old class of anticancer drugs, some of which are perceived as no longer ‘trendy’, new therapeutic approaches based on MTAs continue to be proposed for cancer treatment ^14^. However, due to the development of drug resistance and significant side effects associated with these drugs, there is a constant need for more efficient and specific compounds that target tubulin polymerization. Numerous efforts have been made to develop MTA screening methods. In 2016, the team of John S. Klassen developed an assay of anti-tubulin drugs screening based on catch-and-release electrospray ionization mass spectrometry ^15^. They concluded that the developed assay could be applied for anti-cancer drug screening and for ranking the affinities of compounds to tubulin. Still, they did not apply this new assay to ‘real’ approved chemical libraries. Moreover, the affinities of compounds are not always directly correlated with anti-cancer activity of the molecules. Indeed, there are at least six known binding sites for the small molecules on tubulin, which affect tubulin polymerization differently. Thus, the development of the functional MTA test was still needed, which encouraged the team of Stefano Di Fiore to develop a SNAP-tag-Based screening assay for the analysis of microtubule dynamics and cell cycle progression ^16^. Unfortunately, like the previous assay it was tested only on a small number of molecules, however it follows microtubules functions, making it more appropriate for new MTA screening. The main disadvantage of the proposed assay is that it follows the impact of tested compounds in the cells wherein it is very difficult to separate the direct impact of the molecules on tubulin polymerization from indirect perturbation of cellular cytoskeleton through molecules binding to some other targets that perturb cell cycle and thus impact cytoskeleton. Finally, to the best of our knowledge, until now, there has been no functional high-throughput assay for MTA screening applied to diverse chemical libraries. ^13^

The new nanoDSF screening assay introduced in this study not only overcomes the limitations of previous methods but also introduces additional advantages crucial for accurately determining the mode of action of compounds. Firstly, nanoDSF serves as an *in vitro* functional assay within a simplified environment, enabling the ranking of compounds by their effect on tubulin polymerization, as indicated by shifts in T_poly_. Secondly, nanoDSF incorporates several internal controls that enhance the assay’s reliability. The initial fluorescence measurement confirms the correct tubulin concentration, essential since T_poly_ is concentration dependent. Additionally, this assay enables the determination of tubulin’s T_m_, thereby not only calculating ΔT_m_ for each compound but also ensuring the proper folding state of tubulin at each run. This feature is particularly vital for automated screenings, where maintaining stability of such ‘fragile’ proteins as tubulin over extended periods in plates is a key concern. Furthermore, by employing varying concentrations of compounds, it is feasible to ascertain the apparent association constants of hits with tubulin, based on both the fluorescence signal at a fixed temperature and the denaturation temperature shift. Distinct from previous MTA screening assays, our proposed method has been validated with a chemical library of 1,520 approved compounds, successfully identifying all known tubulin polymerization targets as hits.

### Compounds with highest MTA activity

Through a novel nanoDSF screening assay, we identified approximately 120 compounds with MTA activity within the PCL. Among these, 20 compounds fully inhibited temperature-induced tubulin polymerization under the experimental conditions (Table 1). Some achieved this by directly inhibiting polymerization, while others promoted polymerization at lower temperatures. Notably, recognized MTAs such as Paclitaxel (PTX), Docetaxel, Podophyllotoxin, Colchicine (CLH), Fenbendazole, and Mebendazole were among those that prevented temperature-induced polymerization entirely. Additionally, our study confirmed that artificial estrogens like Hexestrol and Dienestrol, as well as the antiseptic Thiomersal, also impacted tubulin polymerization, consistent with previous reports of their direct effects on tubulin. ^17,18^.

Over half of the strong hits were identified as MTAs for the first time through this screening. Among these, Auranofin (AUF), Ebselen (EBS), and Riboflavin (RBF) had previously been shown to affect the cytoskeleton, despite the absence of direct evidence for tubulin binding. Thus, AUF, an antirheumatic gold complex, inhibits neutrophil activation by markedly reducing the number of centriole-associated microtubules and obstructing phagocytosis in human polymorphonuclear leukocytes, likely through a mechanism that involves microtubule dysregulation ^19,20^. AUF is increasingly recognized as a potential anticancer agent; by serving as an inhibitor of both thioredoxin reductase and proteasome, it induces oxidative stress and triggers apoptosis in models of non-small cell lung cancer (NSCLC)^21^. EBS, an organoselenium compound that mimics glutathione peroxidase, has been explored as a neuroprotectant in ischemia and conditions linked to oxidative stress ^22^. Research demonstrates its ability to destabilize microtubules in skin melanocytes and inhibit tumor growth through the suppression of 6-phosphogluconate dehydrogenase activity ^22–24^. Deficiencies in RBF (vitamin B2) transport are linked to disturbances in microtubule dynamics; however, these disruptions can be alleviated through the administration of Riboflavin in cellular models ^25^. Furthermore, combining Riboflavin with chemotherapy has been suggested as a strategy to reduce side effects and enhance therapeutic outcomes ^26^.

Eight of the identified strong hits were previously unrecognized in their association with tubulin, marking them as novel discoveries. Notably, a family of benzofurans (Benzbromarone (BZB), Benzarone (BZ), and Benziodarone (BZI)) has been demonstrated to directly and effectively modulate tubulin polymerization, aligning with previous findings that BZ inhibits tumor growth *in vitro^27^*. Intriguingly, BZ and BZB, both used in the treatment of gout akin to CLH - a well-known MTA. While BZB is believed to act through uric acid reuptake, CLH disrupts inflammasome assembly at the cytoskeletal level ^28,29^. The revelation that benzofurans may also interact with microtubules introduces an additional dimension to the understanding of their anticancer and anti-inflammatory properties. Among the novel MTAs identified is Aprepitant (APT), a GPCR inhibitor extensively used in chemotherapy to prevent common side effects like nausea. Remarkably, APT is also attributed with antitumor properties of its own. ^30,31^ Bithionol (BTN) and Hexachlorophene (HCP) are fungicides from a class of bridged diphenyl compounds with cytotoxic and antiproliferative action in cancer cell lines ^32,33^. Our findings reveal that several dihydropyridines, approved for managing angina, also interfere with microtubule assembly. Notably, Nifedipine (NFD) and Nisoldipine (NSD) demonstrated the most significant impact on tubulin. While calcium channel blockers like NFD have been previously noted to enhance the sensitivity of drug-resistant cancer cell lines to PTX ^34^, our research marks the first instance of identifying these medications as MTAs.

Collectively, the strong hits identified in this study present compelling cases for drug repurposing, echoing the findings of prior research. Specifically, compounds such as AUF, EBS, APT, BZ, BZB, and HCP exhibit anti-tumor activity *in vitro* ^24,27,33,35–37^. Nifedipine shows potential in reversing drug resistance in cancer cells^34^, while RBF and BTN enhance the effectiveness of existing anticancer drugs ^26,32^. Additionally, BZI and NFD represent modifications of molecules with established antineoplastic properties, further underscoring their potential for repurposing in cancer therapy.

### Structure-activity relationship

#### Carbendazim and benzofuran clusters

Carbendazim derivatives form a significant cluster of compounds with varied MTA activity, ranging from the complete inhibition of tubulin polymerization seen in the presence of Mebendazole (MBZ) and Fenbendazole (FBZ) to a spectrum of high, medium, and low inhibitory effects observed for Albendazole (ABZ, 6.4°C), Nocodazole (NCZ, 6.0°C), Oxfendazole (OFZ, 3.8°C), Methiazole (MTZ, 2.7°C), Parbendazole (PBZ, 1.8°C), Flubendazole (FLU, 1.4°C), and Oxibendazole (OBZ, 1.3°C) (Fig. 3D). While most of these derivatives are utilized as broad-spectrum antihelminthic agents targeting the colchicine site on tubulin, PBZ is employed as an antifungal drug, and only NCZ is used in oncology. However, most exhibit anticancer potential to varying degrees. Specifically, MBZ and ABZ have been highlighted as promising anticancer agents ^38,39^, FBZ has shown moderate antineoplastic activity ^40^, OFZ has been found to inhibit cell growth in NSCLC ^41^, MTZ enhances the efficacy of gemcitabine in pancreatic cancer ^42^, and FBZ has a putative action against triple-negative breast cancer ^43^. Even OBZ, with the lowest inhibitory effect on tubulin polymerization among the MBZ derivatives, has been reported to significantly impede the growth of androgen-independent tumors ^44^. Carbendazim itself, a systemic fungicide with broad-spectrum use, targets tubulin as well ^45^. The most potent inhibitors of tubulin polymerization among its derivatives, MBZ and FBZ, feature a benzene ring attached at the 11^th^ position of the carbendazim structure, connected through a sulfur atom or a carbonyl group. Further analysis reveals that the inhibitory effect on tubulin polymerization, as indicated by changes in ΔT_poly_ for OBZ, PBZ, and ABZ, significantly improves with the substitution of carbon atoms at the first position of the aliphatic chain with sulfur, and to a lesser extent, decreases with substitution by oxygen.

We also identified a compact cluster of three benzofuran derivatives: Benzarone (BZ), Benzbromarone (BZB), and Benziodarone (BZI) (Fig.3A,B). All three compounds exhibited complete inhibition of tubulin polymerization. Considering that the modification of the phenol group with bromine and iodine does not diminish the inhibitory properties of BZ, it suggests that the benzofuran moiety may play a pivotal role in tubulin polymerization inhibition. Supporting this notion, TCZ, featuring a Benzothiophene group - a structure analogous to benzofuran with the oxygen atom replaced by sulfur - exhibits the most pronounced effect on tubulin polymerization within the MCZ cluster. This hypothesis is in line with published data on the impact of benzofuran in its derivatives on tubulin polymerization ^46,47^. While there has been no previous report of these compounds exhibiting MTA activity, BZ has been documented to inhibit the growth of colorectal cancer cells both *in vitro* and *in vivo^27^*. Although the mechanism of BZ’s action was suggested to involve its primary target EYA3 and the inhibition of the EYA3-SIX5-p300 complex, our results suggest the possibility of a direct effect on microtubules as well. This is supported by observations of BZ leading to a dose-dependent decrease in cell proliferation and invasion ^27^, processes fundamentally reliant on MT dynamics. Additionally, BZB has been pinpointed as a potential candidate for drug repositioning in the treatment of lung adenocarcinoma (LUAD) through AI-driven analysis of gene dysregulation ^35^. Our findings lend robust support to these insights, suggesting a potential molecular mechanism behind BZB’s anticancer efficacy.

#### Tricyclic molecules clusters

Several derivatives of Phenothiazine (PTZ) have been identified to exhibit notable MTA activity (Fig. 3). While PTZ itself induces a modest shift in tubulin polymerization temperature (ΔT_poly_) by 1°C, its derivatives - Thiethylperazine (TEP, 1.0°C), Chlorprothixene (CPX, 1.1°C), Toluidine blue (TB, 1.1°C), Perphenazine (PPZ, 1.5°C), Methylene Blue (MB, 1.7°C), Chlorpromazine (CPZ, 1.7°C), Trifluoperazine (TFP, 2.7°C), Flupentixol (FPX, 3.5°C), Fluphenazine (FPh, 3.7°C), and Triflupromazine (TFZ, 6.4°C) - show progressively stronger inhibitory effects (Fig. 3E). These are antipsychotics drugs (except TB and MB) often used in schizophrenia patients and there is epidemiological evidence that has linked lower cancer incidence in schizophrenia patients to long-term medication, highlighting the anticancer potential of antipsychotics^48^. Notably, only TFP and CPZ have been previously recognized for their ability to inhibit microtubule assembly ^49,50^, yet all these derivatives have been either demonstrated or hypothesized to possess anticancer properties, despite their primary classification as central nervous system (CNS) therapeutics targeting dopamine receptors. For instance, PPZ has been highlighted as a potential antitumor agent ^51^, CPZ in oral cancer treatment ^52^, TFP in suppressing colorectal cancer cell models, ^53^, FPX as a potential lung cancer treatment ^54^, FPh in enhancing cancer cell sensitivity to Halaven^55^ and TFZ has been identified as a selective modulator affecting the breast cancer cell cycle^56^. While various mechanisms for the anticancer activity of these drugs have been proposed, our results suggest a direct, shared MTA mechanism among these structurally related molecules. Our data from GBM cell lines and patient-derived tumoroids with varying mutation profiles suggest that certain PTZ derivatives, particularly FPh, have potential for repurposing in glioblastoma therapy.

Additionally, this MTA mechanism might offer an alternative explanation for the pleiotropic effects observed with PTZ derivatives against Gram-negative bacterial persister cells ^57^ and their antitubercular activity ^58,59^. Within the screened PTZ derivative family, there are three pairs of molecules where the substitution of a -Cl group with a -CF_3_ group at the 2^nd^ position (Fig. 3D) markedly enhanced their inhibitory effects on tubulin polymerization. Specifically, for the CPZ and TFZ pair, the ΔT_poly_ escalated from 1.7°C to 6.4°C; for PPZ and FPh, it increased from 1.5°C to 3.7°C; and for prochlorperazine and TFP, it rose from 0.5°C to 2.7°C (Fig. 3D). The critical contribution of the -CF_3_ group to inhibiting tubulin polymerization is also underscored by 1°C higher ΔT_poly_ observed for 2-(Trifluoromethyl) phenothiazine compared to PTZ.

A similar enhancement in tubulin polymerization inhibition resulting from the substitution of a -Cl group with a -CF_3_ group is observed in another pair of molecules, derivatives of thioxanthene. These differ from PTZ derivatives only by the replacement of a nitrogen atom with carbon in the central ring (Fig. 3D). Zuclopenthixol, bearing a -Cl group at the 2^nd^ position, shows no significant effect on tubulin polymerization, whereas Flupentixol (FPX), featuring a -CF_3_ group, induces a 3.5°C shift in T_poly_.

This study also uncovered a group of MTAs among Dibenzosuberane (DBS) derivatives, traditionally recognized as tricyclic antidepressants: Nortriptyline (NTP), Loratadine (LTD), Protriptyline (PTP), Norcyclobenzaprine (nCBP), Opipramol (OPP), Clomipramine (CMP), and Asenapine (ANP). These compounds exhibit a modest effect on tubulin polymerization, with a ΔT_poly_ of approximately 1°C. While none of these were previously recognized for influencing tubulin polymerization, certain members have been noted for their anticancer activities against prostate ^60,61^ or glioblastoma cell lines^62^. Additionally, CMP has been reported to augment the cytotoxicity induced by vinorelbine in human neuroblastoma cancer cells ^63^. The anti-tubulin properties of PTZ, Thioxanthene, and DBS derivatives, which are utilized in CNS treatments and thereby capable of crossing the blood-brain barrier (BBB), position them as promising candidates for repositioning in the treatment of brain tumors.

## Conclusions

In summary, we have developed a novel nanoDSF assay for screening microtubule-targeting agents, compounds widely used in anticancer, antifungal, and antibacterial therapies. Unlike previous assays, our method evaluates both compound binding to tubulin and their impact on tubulin polymerization. We validated this assay using the Prestwick Chemical Library, comprising 1,520 approved compounds, successfully identifying all known MTAs as hits. This approach also uncovered new anti-tubulin drugs among compounds previously associated with cancer cell proliferation inhibition or cytotoxicity, reaffirming tubulin as a critical target for anticancer drug development. Our findings pave the way for drug repositioning of newly discovered MTAs and streamline the search for novel scaffolds in large chemical libraries. We hope to inspire renewed interest in the discovery of anticancer compounds within the MTA class.

## Methods

### Materials

Human glioblastoma cells were from ATCC (Gaithersburg, MD, USA). Compounds used in the cytotoxicity assay were from Prestwick Chemical Library (PCL), MedChemTronica (Bergkällavägen, Sweden) or Sigma (St Louis, MO, USA).

### Tubulin purification

Tubulin was purified from lamb brains by ammonium sulfate fractionation and ion-exchange chromatography and stored in liquid nitrogen as described ^64^. Tubulin concentration was determined at 275 nm with an extinction coefficient of 109 000 M^-1^ cm^-1^ in 6 M guanidine hydrochloride.

### Turbidimetry and differential scanning fluorimetry (DSF) assays

For measurements, aliquots of tubulin were passed through a larger (1 × 10 cm) gravity column of Sephadex G25 equilibrated with 20 mM sodium phosphate buffer, 1 mM EGTA, 10 mM MgCl2, 3.4 M glycerol and 0.1 mM GTP, pH 6.5 (PEMGT buffer) or with 0.8 M PIPES and 0.1 mM GTP, pH 6.9 (PIPES buffer) at 4°C. For MTA binding assays, vinblastine, paclitaxel and colchicine were concentrated up to 400 μM, and tubulin was concentrated at 10 μM. For colchicine, in order to reach the equilibrium of the reaction, we incubated tubulin with colchicine during 90 minutes in the ice. Each capillary was filled with 10 μL of sample. All measurements were performed on nanoDSF Prometheus NT.Plex, from 15°C to 95°C, with optic at 10%, at 1 degree/min.

### NanoDSF Screening

For nanoDSF screening of 1,520 compounds from the FDA-approved list, 50 nanoliter samples in 100% DMSO were placed in 384-cell microplates and stored at −80°C. Prior to nanoDSF measurement, 10 μl of 45 μM tubulin in PEMGT buffer with 0.1 mM GTP was added to each well in a 24-well lane, and mixed thoroughly with pipetting. Final compound concentration in the samples was ∼50 μM in 0.5% DMSO. The plate was centrifuged briefly to avoid bubble formation and samples were trasferred to standard DSF-grade capillaries on a 24-capillary rack. All measurements were performed on nanoDSF Prometheus NT.Plex, from 20°C to 95°C, with optic at 7%, at 1 degree/min.

### Transmission electron microscopy (TEM)

Samples were adsorbed onto 200 mesh, Formvar carbon-coated copper grids, stained with 2% (w/v) uranyl acetate, and blotted to dryness. Grids were observed using a JEOL JEM-1220 transmission electron microscope operated at 80 kV. Magnifications used range from 60’000× to 120’000×. To ensure that microtubules do not disassemble during adsorption, this step was performed in a thermostated room at 37 °C. The same step was performed for grids at 4°C, 55°C and 80°C.

### Cell culture, cytotoxicity and proliferation assays

Glioblastoma cell culture routines, viability and proliferation assays were performed as previously described^65^. U87MG, U87 EGFRvIII, U251MG and T98G cells were maintained in complete MEM media supplemented with 10% FBS, and 2 mM of L-glutamine (Invitrogen, Paris, France). For cytotoxicity assays, cells were counted and plated in 96-well flat-bottom plates (50,000 cells/mL, 5,000 cells per well)). After 24 hours, the cells were treated with increasing concentrations of the MTAs (from 0 to 40 µM) in vehicle solution, containing 0.05% DMSO. All concentrations were done in triplicates. The surviving cells were quantified after 72h by the tetrazolium bromide MTT-assay, according to the manufacturer’s instructions. After cell lysis, the optical density (OD) was measured at 600 nm using Multiskan MS Thermo plate reader (LabSystems, Waltham, MA, USA). Cell viability was expressed as a percentage of survival, using cells treated with vehicle solution as 100%, and the IC50 values were calculated by using Chou and Talalay linearization method^66^. For proliferation assays, cells were counted and plated on PhenoPlate 96-well plates from Perkin-Elmer at 2,000 cells per well. After 24 hours, cells were treated with increasing sublethal concentrations of the MTAs (DMSO was used as control). Pictures were taken every 24 hours using the Perkin Elmer Operetta High Content Imaging System and cells were counted using Harmony imaging and analysis software (Perkin Elkmer). Proliferation was expressed as a percentage.

### Spheroids

U87MG spheroids were cultivated according to the previously described protocol ^67^. Briefly, U87MG cells (1,000 cells per well) were seeded in 96-well U bottom plates containing 100 µl of 20% methylcellulose in the same complete MEM media which was utilized for the cytotoxicity assay (see above). After 72h of cultivation the cells were supplemented with 100 µl of MTA solution, yielding final drug concentrations of 0 to 64 µM in vehicle solution, containing 0.05% DMSO. All concentrations were done in quadruplicates. Spheroid growth was monitored by optical microscopy, using Olympus CKX41 microscope (Olympus LS, Tokyo, Japan). Growth after 6 more days was evaluated by Alamar Blue viability reagent (Invitrogen, Paris, France). Fluorescence reading was performed with POLARstar Omega plate reader (BMG Labtech, Ortenberg, Germany). Cell viability was expressed as a percentage of survival (spheroids treated with vehicle solution were used as 100%) and the IC50 values were calculated as described above.

### Tumoroids

Glioblastoma tumor specimens were collected at Assistance Publique-Hôpitaux de Marseille (AP-HM) within the 4–24 h following surgery. Samples were obtained from the center of biological resources of AP-HM (CRB BB-0033-00097) according to a protocol approved by the local institutional review board and ethics committee (2014-A00585–42). Tumoroids were derived from 3 IDH wild-type glioblastomas as previously described ^12^. Tumoroids were treated with vehicle (DMSO), FPh 17 µM or for PPZ 17.9 µM 72 hours. Ten tumoroids were treated per condition. After treatment, tumoroids were fixed in formol, dehydrated and embedded in paraffin. Immunostainings were performed as previously described ^68^. Slides were then scanned (Nanozoomer 2.0-HT, Hamamatsu Photonics SARL France, Massy, France) and images processed in NDP.view2 software (Hamamatsu). The percentage of Ki67+ cells was determined and the relative quantification of cleaved-caspase-3 positivity is presented according to the intensity and quantity (+, ++, +++).

### Funnel metadynamics

We employed three freely available modern force fields, including Amber19sb. All ligands were parameterized using acpype, with atom point charges derived from *ab initio* 6-31G calculations utilizing psiresp ^69^. Additional parameters were assigned according to the GAFF2 force field. GDP was parameterized in the same manner. The structure of tubulin alpha was modeled based on the coordinates from PDB ID 4o2b ^70^. Protonation states of residues were predicted using PROPKA and manually verified ^71^. The system was then solvated in a triclinic box with periodic boundary conditions using TIP3P water molecules. To neutralize the system and achieve an ionic strength of 0.15 M, Na^+^ and Cl^−^ ions were added. Energy minimization was performed using 5000 steps of the steepest descent method. The equilibration phase comprised seven steps. Initially, a 100 ps NVT simulation was conducted, applying positional restraints of 1000 kJ/(mol·nm²) to the heavy atoms. Temperature coupling was maintained with a velocity rescale thermostat ^72^. This was followed by five rounds of NPT equilibration, each lasting 100 ps, during which restraint strength was gradually reduced: 1000, 500, 200, 100, and 10 kJ/(mol·nm²). Pressure coupling was achieved using a stochastic barostat ^73^.

The funnel metadynamics setup was modeled after the approach described by Raniolo and Limongelli ^74^. A metadynamics potential of 0.5 kJ/mol was applied every 500 steps. Two collective variables were employed. First, the distance between the center of mass (COM) of Colchicine in its binding site and a reference point 20 Å away from its position. This variable projected the ligand along the funnel line. Second, the perpendicular distance of the ligand from this line. A correction to the binding free energy was applied to account for the entropic contribution due to the funnel-shaped restraint, following the equation provided in Raniolo and Limongelli ^74^. The correction factor for the cylinder was calculated as 1.59 kcal/mol. Final binding free energies were reported with error estimates, following the method of Bhakat ^75^, using a statistical analysis window of 1000 ns.

## Declarations

### Ethical Approval

Fresh brain tumor samples were obtained according to a protocol approved by the local institutional review board and ethics committee and conducted according to national regulations (NCT06045065). All the patients supplied written informed consent. All tumor samples were stored in the AP-HM tumor bank (DC-20131781).

### Authors’ contributions

V.E.B.: Conducted nanoDSF and cell survival screening, performed spheroid experiments. R.L.R.: Established the initial nanoDSF setup, conducted ITC and TEM experiments, and prepared the related figures. L.L.: Performed cell proliferation experiments, contributed to data interpretation and analysis. A.S.: Conducted tumoroid experiments and prepared the corresponding figures. R.B.: Contributed to the interpretation of cell survival data and participated in spheroid experiments. D.A.: Purified tubulin for the experiments. C.D.: Managed the chemical library. E.P.: Oversaw chemical management and revise the manuscript. P.R.: Performed SAR analysis. X.M.: Provided supervision for the chemical library. F.D.: Revised the manuscript. A.T.: Analyzed tumoroid experiments and drafted sections of the manuscript. E.T.: Interpreted data from tumoroid experiments and revised the manuscript. A.V.G.: Designed, executed, and analyzed funnel metadynamics experiments, interpreted the data, and drafted sections of the manuscript. P.O.T.: Secured funding, conception and design of the study, supervised the project, performed data analysis and interpretation, prepared Figures, drafted the manuscript, and revised it thoroughly.

### Availability of data and materials

All data available upon request.

