## Supplemental Figure and Tables for "A new nanoDSF approach to anti-tubulin compounds screening revealed novel MTAs among approved drugs"

Supplementary

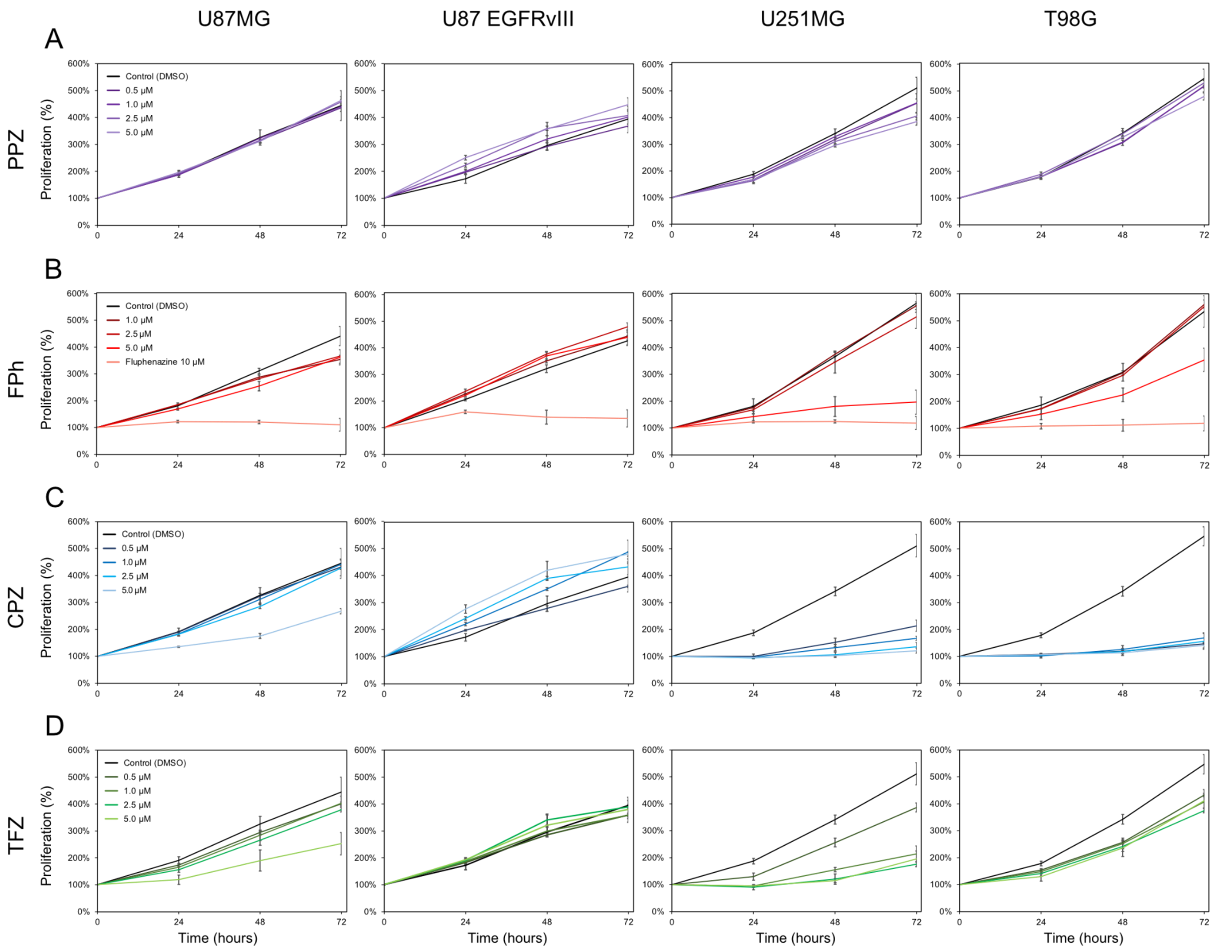


E

| | U87MG | U87 EGFRvIII | U251 | T98G |
|---|---|---|---|---|
| Perphenazine (5 μM) | +5,7% ± 3,9% | +18% ± 3,9% | -30,5% ± 3,4% | -15,0% ± 3,0% |
| Fluphenazine (10 μM) | -96,9% ± 7,2% | -89,3% ± 9,8% | -96,0% ± 3,2% | -95,7% ± 6,3% |

| | U87MG | U87 EGFRvIII | U251 | T98G |
|---|---|---|---|---|
| Chlorpromazine (5 μM) | -51,4% ± 2,8% | +28,8% ± 3% | -94,8% ± 2,3% | -90,7% ± 1,1% |
| Triflupromazine (5 μM) | -55,7% ± 12% | -4,1% ± 12,8% | -76,6% ± 7,0% | -30,2% ± 9,1% |

Figure 1S. (A-D) Effects of escalating concentrations of PPZ, FPh, CPZ, and TFZ on the proliferation of different glioblastoma cell lines U87MG, U87 EGFRvIII, U251MG, and T98G. (E) Increases in the proliferation of U87MG, U87 EGFRvIII, U251MG, and T98G glioblastoma cell lines at maximal sublethal concentration of PPZ, FPh, CPZ, and TFZ.

Table S1. Phenothiazine, Thioxanthene, and Dibenzosuberane clusters

| Name | Code | $\Delta T_{poly}$,°C | $IC_{50}$, µM |
|---|---|---|---|
| **Phenothiazine group** | | | |
| Phenothiazine | PTZ | 1.0 | 60.2±9.2 |
| Thiethylperazine | TEP | 1.0 | 8.4±0.2 |
| Perphenazine | PPZ | 1.5 | 9.8±0.4 |
| Chlorpromazine | CPZ | 1.7 | 4.5±0.5 |
| Trifluoperazine | TFP | 2.7 | 17.3±3.2 |
| Fluphenazine | FPh | 3.7 | 7.5±0.5 |
| Trifluopromazine | TFZ | 6.4 | 6.5±0.6 |
| **Thioxanthene group** | | | |
| Chlorprothixene | CPX | 1.1 | 27.6±2.2 |
| Flupentixol | FPX | 3.5 | 9.8±1.8 |
| **Dibenzosuberane group** | | | |
| Opipramol | OPP | 1.0 | n/d |
| Protriptyline hydrochloride | PTP | 1.1 | 17.7±1.5 |
| Asenapine | ANP | 1.1 | 22.5±4.0 |
| Clomipramine | CMP | 1.6 | 19.6±0.6 |
| **Other tricyclic compounds** | | | |
| Toluidine blue | TB | 1.1 | 6.3±0.3 |
| Methylene Blue | MB | 1.7 | 64.2±9.9 |
| Riboflavin | RBF | n/a | ≫60 |

n/a - not applicable; n/d - not determined.

Table S2. Carbendazim and Coumarone clusters

| Name | Code | $\Delta T_{poly}$,°C | $IC_{50}$, µM |
|---|---|---|---|
| **Carbendazim group** | | | |
| Fenbendazole | FBZ | n/a | 0.2±0.0 |
| Mebendazole | MBZ | n/a | ≪0.1 |
| Astemizole | ASZ | 1.1 | 1.7±0.1 |
| Oxibendazole | OBZ | 1.3 | 1.8±0.3 |
| Flubendazole | FLU | 1.4 | 0.6±0.0 |
| Parbendazole | PBZ | 1.8 | ≪0.1 |
| Triclabendazole | TCZ | 2.2 | 15.1±0.9 |
| Methiazole | MTZ | 2.7 | n/d |
| Oxfendazole | OFZ | 3.8 | 17.0±1.0 |
| Nocodazole | NCZ | 6.0 | <0.1 |
| Albendazole | ABZ | 6.4 | 0.3±0.0 |
| **Coumarone group** | | | |
| Benzarone | BZ | n/a | 7.4±0.7 |
| Benzbromarone | BZB | n/a | 62.1±2.8 |
| Benziodarone | BZI | n/a | n/d |

n/a - not applicable; n/d - not determined.

Table S3. Miconazole cluster

| Name | Code | $\Delta T_{poly}$,°C | $IC_{50}$, µM |
|---|---|---|---|
| Isoconazole | ICZ | 1.9 | 33.1±1.8 |
| Miconazole | MCZ | 2.3 | 12.9±0.7 |
| Sertaconazole nitrate | STZ | 3.8 | 10.7±1.3 |
| Tioconazole | TOZ | 4.3 | 18.3±1.4 |

Table S4. Nifedipine cluster

| Name | Code | $\Delta T_{poly}$,°C | $IC_{50}$, µM |
|---|---|---|---|
| Nifedipine | NFD | n/a | 40.4±6.6 |
| Nisoldipine | NSD | n/a | 25.3±1.6 |
| Felodipine | FDP | 1.2 | 13.7±4.0 |
| Nitrendipine | NTD | 1.4 | 36.8±3.3 |

n/a - not applicable.

Table S5. Stilbenoids clusters

| Name | Code | $\Delta T_{poly}$, °C | $IC_{50}$, μM |
|---|---|---|---|
| **Stilbestrol group** | | | |
| Dienestrol | DE | n/a | 23.6±1.0 |
| Hexestrol | HXS | n/a | 20.2±0.5 |
| Diethylstilbestrol | DES | 2.7 | n/d |
| **Tamoxifen group** | | | |
| Toremifene | TMF | 1.9 | 13.3±1.9 |
| (Z,E) Clomiphene citrate | CMX | 2.6 | 5.0±1.0 |
| Tamoxifen citrate | TMX | 7.0 | n/d |

n/a - not applicable; n/d - not determined.

Table S6. Steroids cluster

| Name | Code | $\Delta T_{poly}$,°C | $IC_{50}$, μM |
|---|---|---|---|
| Methandrostenolone | MAS | 1.4 | 58.7±5.1 |
| 17β-estradiol | ED | 1.5 | 30.7±1.9 |
| Norgestimate | NGS | 2.8 | ≫60 |
| Ethinylestradiol | EES | 3.9 | 15.7±0.8 |
| Deflazacort | DFC | −1.3 | ≫60 |
| Gestodene | GD | −2.8 | n/d |

n/d - not determined.

Table S7. Diphenyls clusters

| Name | Code | $\Delta T_{poly}$,°C | $IC_{50}$, μM |
|---|---|---|---|
| **Benzophenone group** | | | |
| Oxybenzone | OXY | 1.5 | 14.4±7.0 |
| Dioxybenzone | DOB | 1.8 | ≫60 |
| **Diphenylmethane group** | | | |
| Hexachlorophene | HCP | n/a | 5.1±0.7 |
| Lidoflazine | LFZ | 1.1 | n/d |
| Diclazuril | DCZ | 1.7 | 48.9±6.0 |
| Dichlorophen | DCP | 2.7 | 7.7±1.2 |
| Mitotane | MTT | 2.7 | 23.7±2.7 |
| **Diphenyl ethers group** | | | |
| Triclosan | TCL | 2.9 | 11.6±0.5 |
| Liothyronine | LTY | −2.7 | ≫60 |
| (L)-Thyroxine | TYX | −2.6 | ≫60 |
| **Other diphenyl compounds** | | | |
| Bithionol | BTN | n/a | ≫60 |
| Meclofenamic acid | MFA | 2.9 | n/d |

n/a - not applicable; n/d - not determined.

Table S8. Non-clustered compounds

| Name | Code | $\Delta T_{poly}$,°C | $IC_{50}$, µM |
|---|---|---|---|
| **Known MTAs** | | | |
| Thimerosal | TMS | n/a | 2.1±0.1 |
| Ethacrynic acid | ECA | 1.3 | ≫60 |
| Griseofulvin | GSF | 2.0 | 24.4±3.7 |
| Phenolphthalein | PP | 3.6 | ≫60 |
| Disulfiram | DFM | −6.5 | 12.0±1.6 |
| **New MTAs with known anti-cancer activity** | | | |
| Aprepitant | APT | n/a | 41.8±31.8 |
| Auranofin | AUF | n/a | 0.9±0.6 |
| Ebselen | EBS | n/a | 50.9±32.7 |
| Flunarizine | FNZ | 1.0 | 13.3±1.2 |
| Acemetacin | AMT | 1.1 | ≫60 |
| Fluoxetine | FX | 1.1 | 26.5±4.1 |
| Flurbiprofen | FBF | 1.2 | ≫60 |
| Troglitazone | TGZ | 1.3 | 39.5±1.0 |
| Propofol | PPF | 1.4 | 26.6±2.5 |
| Mefloquine | MFQ | 1.5 | 11.6±1.2 |
| Clotrimazole | CTM | 1.6 | 6.1±0.6 |
| Sertindole | STD | 2.2 | 6.7±0.2 |
| Efavirenz | EFV | 5.6 | 26.7±2.6 |
| Tizanidine | TZN | −1.3 | n/d |
| Haloprogin | HPG | −1.4 | n/d |
| Mesalamine | MSL | −1.6 | ≫60 |
| Artesunate | ASN | −1.6 | 24.0±4.1 |
| Nafamostat | NFM | −2.8 | 45.2±6.1 |
| **New MTAs with unknown effect on cancer cells** | | | |
| Merbromin | MBR | 2.4 | ≫60 |
| Dihydrostreptomycin | DHS | 1.3 | ≫60 |
| Moguisteine | MGS | −1.0 | n/d |
| Clorsulon | CLS | −1.1 | 66.8±13.3 |
| Pramipexole | PPX | −1.2 | n/d |
| Paroxypropione | POP | −2.5 | n/d |
| Colistin | CST | −3.0 | ≫60 |

n/a - not applicable; n/d - not determined.
